# α-Linolenic acid modulates phagocytosis of extracellular Tau and induces microglial migration

**DOI:** 10.1101/2020.04.15.042143

**Authors:** Smita Eknath Desale, Subashchandrabose Chinnathambi

**Affiliations:** Neurobiology Group, Division of Biochemical Sciences, CSIR-National Chemical Laboratory, Dr. Homi Bhabha Road, 411008 Pune, India; Academy of Scientific and Innovative Research (AcSIR), 411008 Pune, India

**Author notes:** To whom correspondence should be addressed: Prof. Subashchandrabose Chinnathambi, Neurobiology group, Division of Biochemical Sciences, CSIR-National Chemical Laboratory (CSIR-NCL), Dr. Homi Bhabha Road, 411008 Pune, India.

**Keywords:** Fatty acids, α-Linolenic acid, phagocytosis, MTOC repolarization, Tauopathy

## Abstract

The seeding effect of extracellular Tau species is an emerging aspect to study the Tauopathies in Alzheimer’s disease. Tau seeds enhance the propagation of disease along with its contribution to microglia-mediated inflammation. Omega-3 fatty acids are known to exert the anti-inflammatory property to microglia by modulating cell membrane compositions that influence various receptors expression and signaling cascade. The immunomodulatory function of omega-3 fatty acids exerts anti-inflammatory properties to microglia. Owing to the imparted anti-inflammatory nature enhance phagocytosis and increased migration property has been observed in microglia. The increased phagocytosis of extracellular Tau monomer and aggregates has been observed upon ALA exposure to microglia cells. The intracellular degradation of internalized Tau species was targeted by early and late endosomal markers Rab5 and Rab7. The increased levels of LAMP-2A and colocalization with internalized Tau indicated the degradation *via* lysosome. These results indicate the degradation of internalized Tau species in the presence of ALA instead of getting accumulated in the cell. The enhanced migratory ability of microglia in the presence of ALA induces the MTOC repolarization. Tau seeds greatly contribute to the spread of disease, one way to reduce the spreading is to reduce the presence of extracellular Tau seed. Microglia could be influenced to reduce extracellular Tau seed with dietary fatty acids. Our results suggest that dietary fatty acids ALA significantly enhances phagocytosis and intracellular degradation of internalized Tau. Enhanced migration supports the phagocytosis process. Our approach provides insights into the beneficial role of ALA as an anti-inflammatory dietary supplement to treat AD.

## Introduction

In central nervous system (CNS), embryonic mesoderm-derived microglia are the major group of resident immune cells, which consists of 20% of the total glial population. In a normal physiological condition, microglia displays ramified morphology having long branched cellular processes, which senses the tissue damage, pathogenic invasions, *etc*. ^1^. The surveillant stage of microglia is maintained by neuronal and astrocytes-derived factors ^2^. On external stimuli, microglia become activated, and the ramified morphology changes to amoeboid morphology. Microglia are either classically activated to give a proinflammatory response or follow alternative activation to show anti-inflammatory response. Alzheimer’s disease (AD), which is a progressive neurodegenerative disease shows a predominance of inflammatory microglia. Gliosis in AD pathology indicates abnormal morphology, excessive activation of microglia, and astrocytes. Neuroinflammation acts as a key triggering process in AD where Amyloid-β and neurofibrillary tangles of Tau are found to surrounded by microglia^3^. The presence of excessive pro-inflammatory cytokines IL-1β, TNF-α, IFN-γ, and accumulation of aggregated protein’s proinflammatory condition of microglia favors and hampers the phagocytic nature of microglia^4^. The phagocytic state of microglia is regulated by the expression of receptors on the cell surface, membrane fluidity, downstream signaling, and rearrangement of the actin network. Phagocytic ability of microglia is under influence of various environmental factors, lipids is one of them^5^. Dietary lipids affect the brain extensively since fatty acids are building blocks of the brain. Dietary fatty acids importantly polyunsaturated fatty acids (PUFAs) including omega-3 fatty acids Docosahexaenoic acid (DHA-22 n3:6), Eicosapentaenoic acid (EPA-20 n3:5) and α-Linolenic acid (ALA 18 n3:3) have beneficial effects on the brain. Omega-3 fatty acids enhance fluidity of cell membrane by incorporating long-chain fatty acids into phospholipids of cell membrane. The increased fluidity of the cell membrane holds the extent of receptor expression on the cell surface and their downstream signaling^6^. DHA and EPA are either taken up by dietary lipids or synthesized by α-Linolenic acid in the body. DHA and EPA are the main regulators of lipid mediators that drive the resolution phase by suppressing the inflammatory response and helps to restore the homeostasis ^6, 7^.

Microtubule-associated protein-Tau, which forms neurofibrillary tangles (NFTs) in the neurons is considered one of the major consequences of AD along with extracellular amyloid-β plaques ^8^. Glial activation acts as a major cause to drive the pathology-associated with AD. The establishment of Tau as a factor of neurotoxicity and neuroinflammation is still a matter of debate, but recently accepted the concept of Tau as a prion-like protein that supports this hypothesis ^9, 10^. The spreading of Tau and its ability to cause template-dependent deformation in the healthy neuron can be targeted^11, 12^. Omega-3 fatty acids are found to implement the suppression of neuroinflammation and triggers polarization of microglia^13^. Omega-3 fatty acids elevate the resolution phase and mediate tissue repair, healing, and clearing of debris and maintain homeostasis by microglia. Enhanced phagocytic nature of microglial cells due to exposure of Omega-3 fatty acids could act as a therapeutic strategy to minimize the spreading of Tau in Tauopathies to reduce the propagation of disease ^11, 14^. In the process of phagocytosis, Rab proteins especially Rab5, 7 play an important role in intracellular vesicle trafficking and mediates the endocytic pathways. Rab5 is associated with early endosomal marker whereas; Rab7 identifies late endosomes in the phagocytosis process. Hence, to study the internalization and the subsequent degradation of Tau, Rab 5, Rab 7, and their transition can help to provide the insights of the process ^15, 16^. The final step of phagocytosis involves a fusion of late endosomes with lysosome to form phagolysosome as a microcidal compartment. The fusion of late-endosome involves LAMP-lysosome-associated membrane proteins and other luminal proteases ^17, 18^. The high acidic pH (pH-4.5) along with other hydrolytic enzymes (cathepsins, proteases, lysozymes) ensures the elimination and degradation of microorganisms ^19, 20^. Microglia activation leads to polarization and migrates in a particular direction, depending upon the directional clues. Cytokine and chemokine are a response to play an important role in migration and polarization of microglia such as CX3CL1-CX3CR1, IL-4, CCR5, CCR3, and CCR1 mediated signalling 21. The polarized state of microglia is maintained by the cytoskeletal network where actin provides directional sensing and microtubule dynamics for the mechanical strength to move cell forward ^22^. In the process of phagocytic cup formation along with actin network Iba-1(ionized calcium adapted molecule-1) protein of microglia plays an important role. Iba-1 is also reported to have a key role in the function of activated microglia^23, 24^.

In this study we have studied α-Linolenic acid as a precursor for the DHA and EPA, to increase the phagocytic capacity of microglia by enhancing the phagocytosis of extracellular propagating Tau; eventually reducing their spreading. The downstream degradation of internalized Tau was assessed by endosomal transition with Rab5, Rab7 endosomal markers and LAMP-2A as a lysosomal marker to track the degradation of internalized Tau. The migration of microglia has been studied as one of the key properties of the alternative anti-inflammatory phenotype of microglia. The migration profile of microglia was studied with wound scratch assay, reorientation of the microtubule-organizing center along with nuclear centrosomal (NC) axis, and actin rearrangement as morphological hallmarks of activated microglia. We have analyzed the enhancement of phagocytosis after ALA exposure and the degradation was confirmed with the localization with Rab 5, 7, and LAMP-2A. The ALA also improves the migration profile of microglia that aids to the phagocytosis.

## Results

### Tau aggregation in the presence of α-Linolenic acid (ALA)

α-Linolenic acid (ALA) is an essential omega-3 fatty acid, which is a precursor of Docosahexaenoic acid (DHA) and Eicosapentaenoic acid (EPA) ^27^. The role of Omega-3 fatty acids in cardiovascular diseases is well-studied but its role in neuroprotection is needs to be studied ^28, 29^. In this study, we aim to understand the role of ALA on the function of microglia and its effect on extracellular Tau in Alzheimer’s disease. Tau, a natively unfolded protein stabilizes microtubules in neuron and other CNS cells. The longest isoform of Tau has 441 amino acids with two inserts, proline-rich domain and four imperfect repeat regions (Fig 1a). The positive charge of the repeat region of Tau facilitates the binding of anionic free fatty acids ^30^. The beneficial effect of ALA as a potent anti-inflammatory agent and a precursor of other omega-3 fatty acids DHA, EPA provides therapeutic strategy in AD. In this study, we explored the neuroprotective anti-inflammatory role of ALA on exposure to microglia and its effect on phagocytosis of extracellular Tau (Fig. 1b). ALA is a polyunsaturated omega-3 fatty acid (18:3) having three double bonds in its structure and it is a precursor of DHA and EPA. For the preparation of ALA was dissolved in 100% ethanol and then solubilized at 50°c, they produce vesicles-like structure, which has been shown in transmission microscopy (TEM) images (Fig. 1 c, d). Due to high hydrophilic nature, high net positive charge and lack of hydrophobic residues accounts for the natively unfolded nature of Tau. This flexible structure of Tau due to unfolded nature aids for microtubule-binding and stability. The highly soluble form of Tau can be induced to aggregate in the presence of polyanionic agents such as heparin, which neutralize net positive charge *in vitro*. The hTau40 aggegates produced *in vitro* with heparin and their characterization with different biochemical assays are enlisted by diagrammatic representation. Free fatty acids such as arachidonic acid induce spontaneous self-assembly of Tau protein to form aggregates in dose-dependent manner^30^. *In vitro* aggregation of hTau40 in presence of heparin was confirmed with ThS fluorescence for time period of 120 hours, SDS PAGE analysis and TEM for visualization of aggregated Tau fibrils (Fig 1 e, f, and g). The confirmation for the aggregates formation in presence of Tau was carried out with the circular dichroism spectroscopy (CD). The native random coil nature of Tau changes to β-sheet conformation on formation of aggregates which can be detected with the shift in absorbance in CD data (Fig. 1 h).

**Figure 1.**
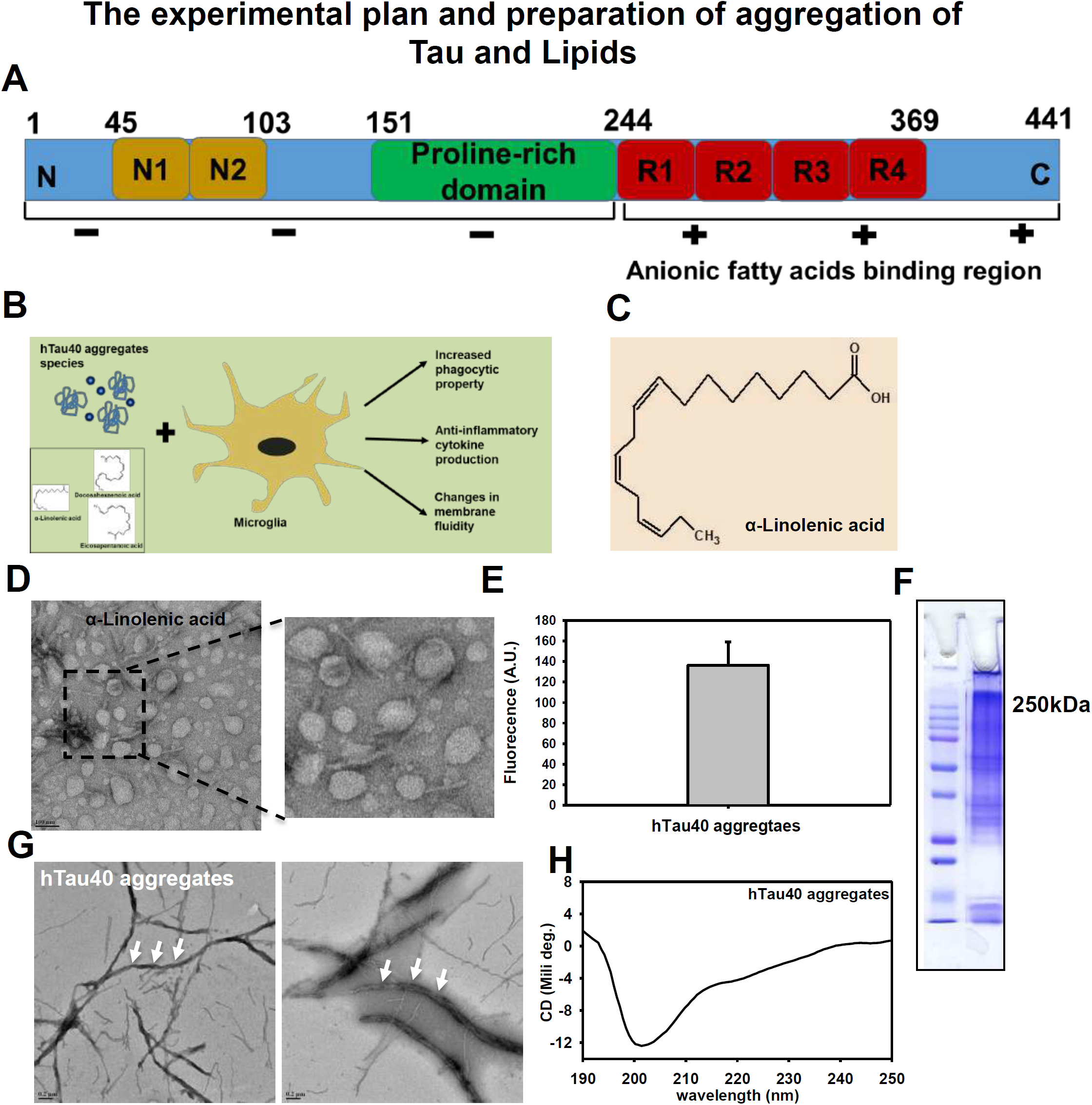
Biochemical characterization of hTau40 aggregates. Experimental approach and biochemical characterization of ALA: A) Tau structure bar diagram showing domains of hTau40 having 441 amino acid sequence and specified with the distribution of net charge domain vise. The fatty acid binding region is indicated at repeat region of Tau structure. B) The proposed hypothesis for the effect of ALA and hTau40 species on microglia, ALA changes the membrane composition of microglia and enhances anti-inflammatory phenotype with increased phagocytic capacity and also modulates membrane fluidity; we propose that increased phagocytic ability would clear the extracellular Tau species. C) Chain structure of α-Linolenic acid (ALA) (18 n: 3). D) ALA was dissolved in 100% ethanol and solubilized at 50 degrees for 2 hours. The microscopic observation of ALA vesicles was done by transmission electron microscopy for the morphological analysis. The enlarged area showing zoomed images of vesicles; scale bar is 100 nm E) ThS fluorescence assay, to observe the aggregation propensity of hTau40 at 120 hours time points in presence of heparin *in vitro*. F) SDS PAGE analysis of hTau40 aggregates. The marking of 250 kDa shows higher order bands corresponding to aggregates. G) TEM analysis of hTau40 aggregates after 120 hours. H) CD analysis to study the conformation changes of Tau on aggregation from random coiled to β-sheet structure. The spectra was analysed between 250-190 nm range.

### Internalization of extracellular Tau in presence of ALA in microglia

Long chain polyunsaturated Omega-3 fatty acids are integral part of membrane phospholipid^31^. Incorporation of long chain fatty acid in microglia cell membrane increases the fluidity of membrane and hence enhances anti-inflammatory phenotype ^5^. The intrinsic phagocytosis property of microglia enhances as exposed to long chain polyunsaturated fatty acids (PUFAs) since in case of microglia PUFAs exerts anti-inflammation properties and suppresses pro-inflammatory properties. We expose microglia cells (N9) with 40 µM ALA for 24 hours and checked for the internalization of extracellular Tau monomer and aggregates (Fig. 2a). N9 cells were treated with 40 µM ALA as a control, 1 µM Tau monomer, aggregates and their respective treatment with ALA. Immunofluorescence staining was performed to study the internalization of Tau (red) in Iba-1 (green) positive microglia cells since Iba-1 is marker for microglia and involves in membrane-ruffling and phagocytosis ^23^. Phagocytosis of extracellular Tau has increased in both Tau monomer and aggregates exposure in presence of ALA (Fig. 2b). The internalization of extracellular aggregates observed to be increased as compared to extracellular monomer, thus indicates that ALA enhances phagocytosis ability of N9 cells. 3-D view of immunofluorescence images indicates the presence of internalized Tau. The insight representation of single cell of 3-D immunofluorescence images indicated that the internalized Tau shown with the white arrow marks. Intracellular intensity of internalized Tau was quantified in fluorescence images, significant increase in internalization quantified as an intensity/µm^2^ area was observed in cells treated with ALA as compared to control (no treatment) cells (P<0.001) (Fig. 2c). ALA exposure increased the intrinsic phagocytic capacity of microglia in monomer and aggregates by 68 and 75% with P< 0.05, 0.01 respectively (Fig 2 d). Supplementary figure 1a (Fig. S1 a) incorporates the individual panel for all the filters given in the merge images for better understanding of morphology and immunofluorescence staining as Tau (red), Iba-1 (green), DAPI (blue) and DIC (Differential interference contrast). Supplementary figure 1b (fig. S1 b) shows the orthogonal view indicating xz and yz axis for the better understanding of localization of Tau.

**Figure 2.**
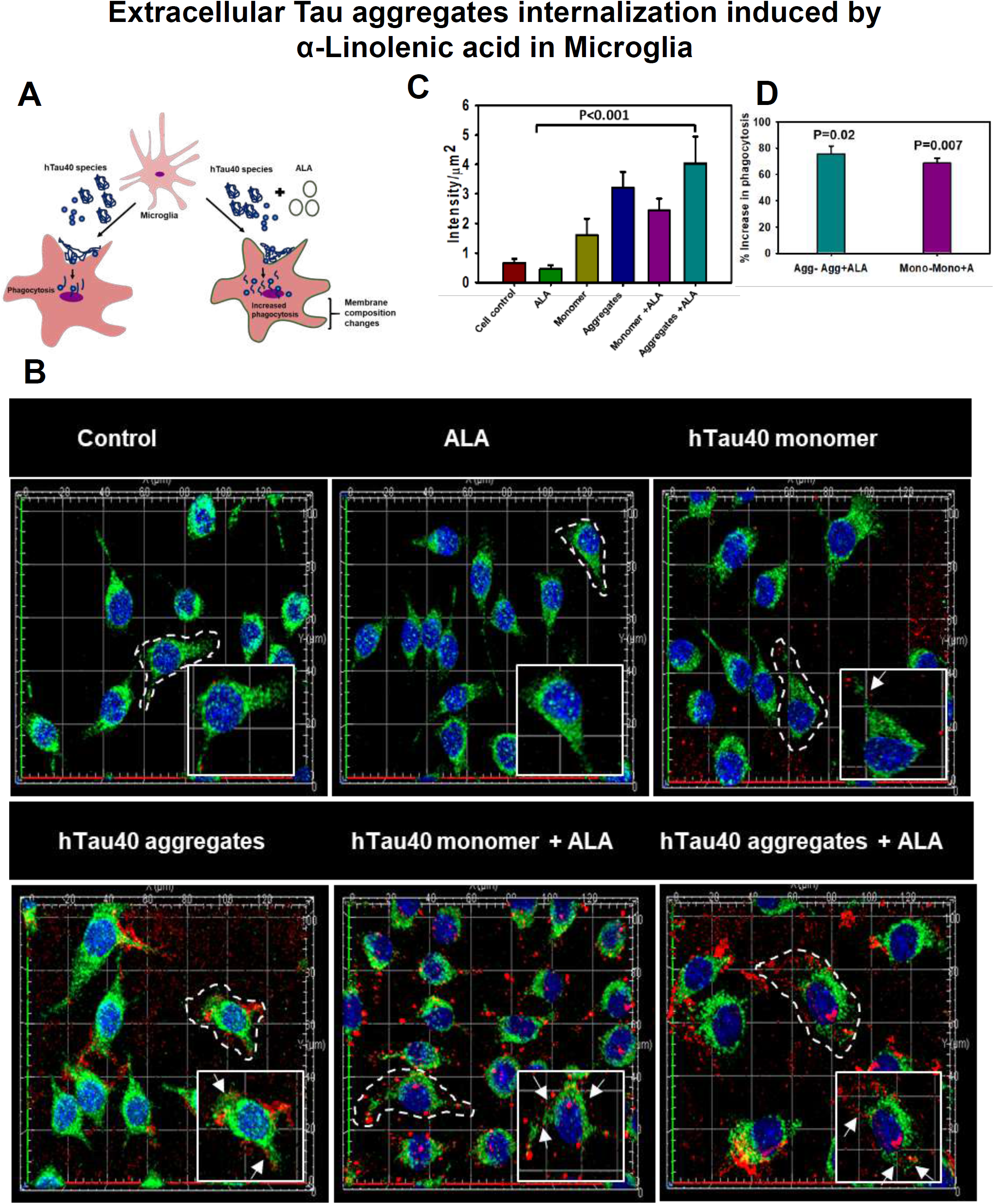
Extracellular Tau aggregates internalization, induced by α-Linolenic acid in Microglia. Internalization of hTau40 recombinant Tau in Iba-1 positive microglia. A) ALA is one of the omega-3 fatty acids which has major role on cell membrane modulation of microglia, increased ALA in diet induces cell membrane changes that enhances the anti-inflammatory phenotype of microglia, we hypothesize that increased phagocytic ability of microglia may clear the extracellular Tau species which are responsible for Tau seeding across the healthy cells. B) Cells were incubated with hTau40 aggregates species, hTau40 monomer species (1µM) alone and along with the α-Linolenic acid (40µM) for 24 hours at 37°c as control 40 µM ALA alone and cell control (without treatment) was kept for comparison. The cells were fixed after 24 hours and stained with anti-Iba-1 antibody (green) and T46 Tau antibody (red) and observe by fluorescence microscopy, scale bar is 20µm. The enlarged image showing the zoomed areas of microglia with internalized Tau marked with white arrow. C) 3D view of internalized recombinant hTau40 in Iba-1 positive microglia. 3D view helps to show the localization of internalized Tau in microglia. The images were taken with the Zeiss fluorescent microscope with Apotome 2.0. E) Quantification of internalized Tau intensity per unit square area of microglia cells; showing extent of internalization Tau in microglia cells which is highly significant P<0.001 compared with cell control. D) Percentage increase in phagocytosis of hTau40 in microglia after ALA exposure to cells; percentage increase in aggregates to aggregates with ALA exposed groups and monomer to monomer with ALA groups calculated from the intracellular intensity of Tau amongst the groups, significance is P= 0.02, 0.007 respectively.

### Effect of ALA on endosomal trafficking of internalized Tau and its degradation pathway

The phagosomes after internalization is subjected to lysosome-mediated degradation *via* endosomal maturation process in phagocytosis ^32, 33^ The phagosomes after internalization fuses with endosomal compartments, by endosomal markers Rab5, Rab7, where maturation of endosome occurs and finally it fuses with lysosomes for degradation of internalized microorganisms in immune cells. We expected the colocalization of internalized Tau with endosome compartment since the endosomal maturation is followed to degradation pathway. We studied downstream early and late endosomal markers Rab5, Rab7 and LAMP-2A respectively for the colocalization with internalized Tau (Fig. 3a). The immunofluorescence images of Tau and endosomal, lysosomal markers after 24 hours of exposure with extracellular Tau monomer, aggregates and ALA showed the levels of endosomal and lysosomal markers in the cell and the colocalization with internalized Tau represented with white arrow marks in images. (Fig. 3b, 4a). The zoomed images shows the area of colocalization of internalized Tau with Rab5 and Rab7 in microglia (Fig. 3b, 4a). The intracellular intensity per unit area of Rab5 and Rab7 was estimated.

**Figure 3.**
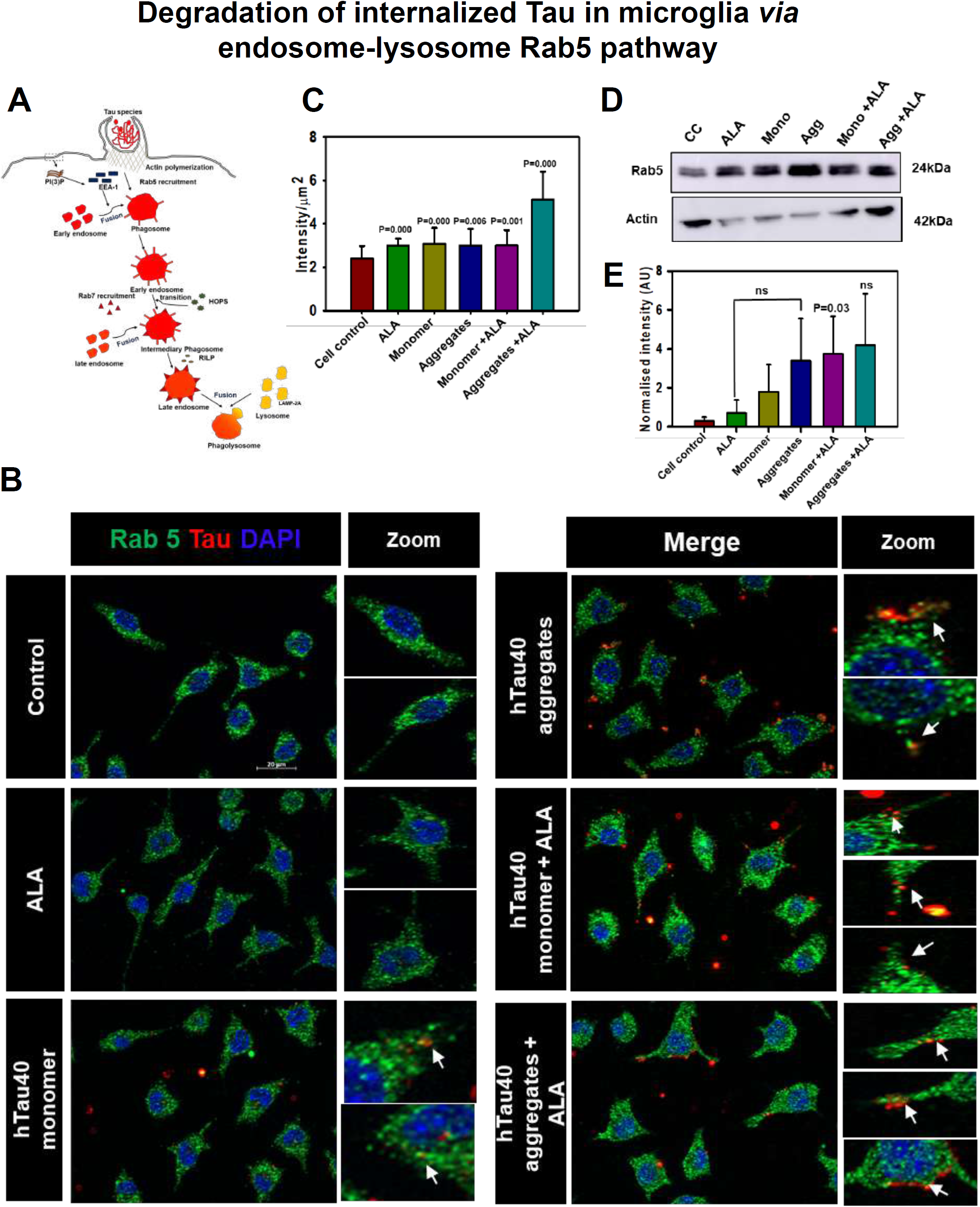
Degradation of internalized Tau in microglia via endosome-lysosome pathway. A microglia cell were exposed to hTau40 monomer and aggregates in presence and absence of ALA and observed for the levels of Rab 5 (green), and Tau (red) by fluorescence microscopy. The degradation of internalized Tau was studied with the early endosomal marker and late endosomal markers. A) The internalized Tau follows the phagocytosis pathway and finally degrades via lysosome-mediated degradation. In the pathway the maturation of phagocytic vesicle take place that can be marked with the early endosomal marker Rab 5 and late endosomal marker Rab 7. We observed the colocalization of internalized Tau with endosomal markers to trace the degradation of internalized Tau. B) The fluorescence microscopy images indicates the levels of endosomal markers and their colocalization with internalized Tau, the zoomed area indicates the colocalized positions inside the cell, the white arrow marks indicates colocalization. C) The intensity analysis of endosomal markers was carried out and plotted as intensity per unit sq. area; significance is P<0.05. D) Expression analysis of early endosomal marker was observed by western blot after various treatments of hTau40 monomer, aggregates and ALA after 24 hours.

**Figure 4.**
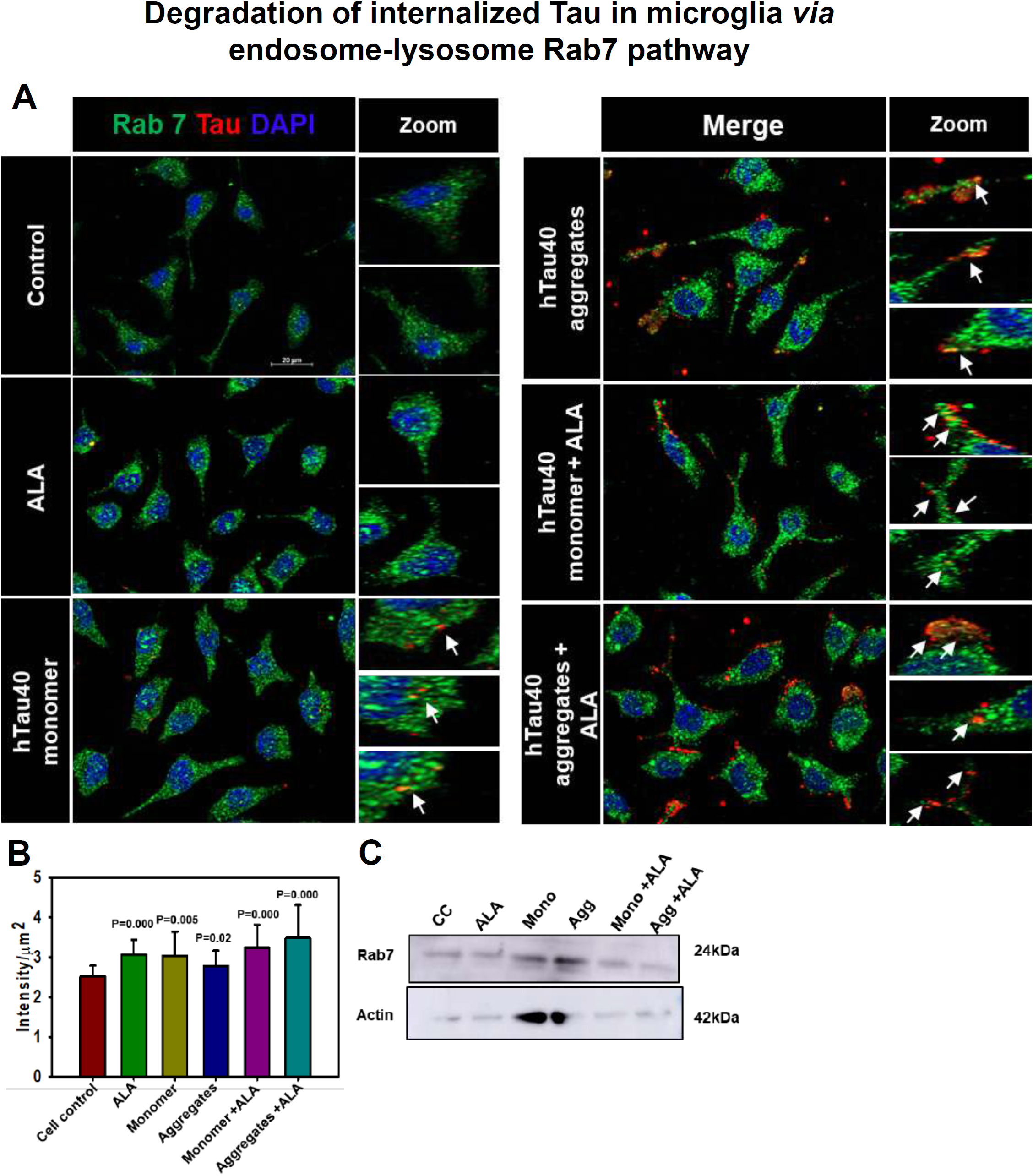
Degradation of internalized Tau in microglia via endosome-lysosme pathway. Fate of internalized Tau was observed with the help of late endosomal marker Rab 7 by fluorescence microscopy. A) The colocalization of internalized Tau was observed with late endosomal marker Rab 7 and the zoomed area indicates the area of colocalization inside the cell. B) The intensity analysis of Rab 7 plotted as intensity per unit sq. area, significance is P<0.05. C) Expression profile of Rab 7 by western blot after treatment conditions.

The expression of Rab5 was found to increased significantly in case of aggregates with ALA treated cells (p<0.001), whereas Rab7 showed increased levels of protein with ALA treatment in both monomer and aggregates treated cells as compared to other control groups (P<0.001) (Fig. 3c, 4b). Expression of Rab5 and 7 was also checked with western blot, aggregates treated groups, where we found to increase both Rab5 and 7-protein levels (Fig. 3d, 4c). The increased levels of Rab 5 and 7 in case of ALA treated cells in both monomer and aggregates shows cells are undergoing more of phagocytosis and the internalizing Tau is channelizing towards degradation pathway. The mature late-endosome containg target then fuses with the lysosome. The high pH in lysosome compartment and other hyderolytic enzymes containg proteases, lipases, lysozymes, cathepsins induce degradation of internalized targets ^34^.

### Effect of ALA on lysosome-mediated degradation of internalized Tau

Further to understand lysosome-mediated degradation, N9 cells were treated with extracellular Tau monoer, aggregates and ALA and the cells were stained for immunofluorescence analysis with Tau (red) and LAMP-2A(green) post treatment. The levels of LAMP-2A and its colocalization internalized Tau indicated through immunofluorescence analysis (Fig. 5a). The 3-D representation of immunofluorescence images indicated colocalization of Tau and LAMP-2A, zoom image panel indicates colocalization spotted with white arrow marks (Fig. 5a). The intracellular intensity of LAMP-2A was calculated, which does not show changes in intensity (Fig. 5b). the levels of LAMP-2A by western blot was analysed, the ALA treated N9 cells showed increase in levels in both Tau monomer and aggregates exposed cells (Fig. 5c, d). For the better understanding of internalized Tau orthogonal view of immunofluorescence images was provided, the x and y axis of the images shows localization of Tau in cell (Fig. S2).

**Figure 5.**
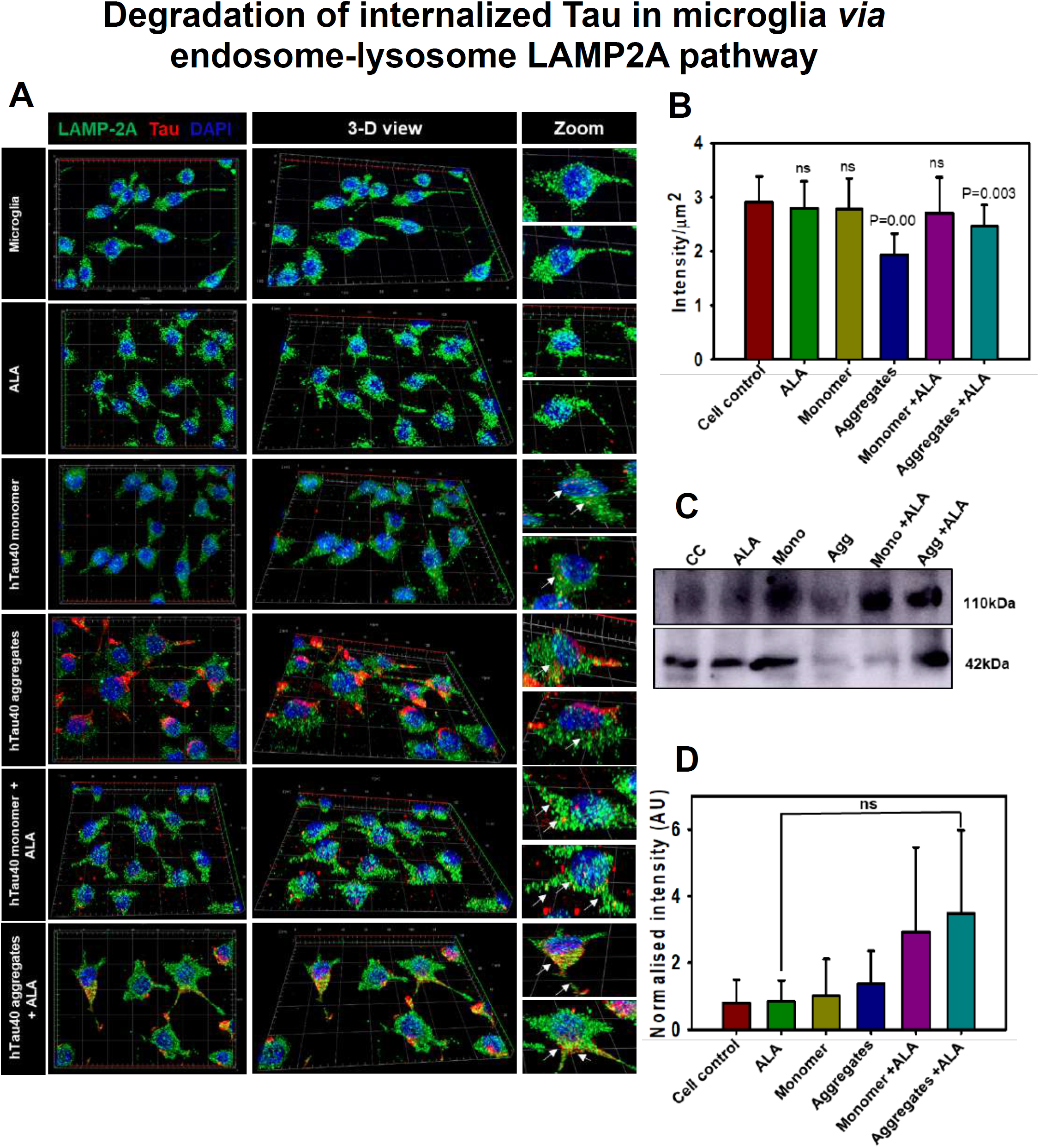
Degradation of internalized Tau in microglia via endosome-lysosome pathway. Last step of degaradation pathway includes fusion of late endosome with lysosome studied by fluorescence microscopy. A) Internalization of extracellular Tau (red) studied with its colocalization with LAMP-2A (green) after 24 hours of ALA and extracellular Tau exposure. The 3D images showing levels of LAMP-2A and its colocalization with Tau. Zoom panel shows enlarged area from the 3D image showing colocalization of Tau and LAMP-2A. B) Intracellular intensity of LAMP-2A was calculated from immunofluorescence images and plotted as intensity per unit sq. area. C) Levels of LAMP-2A was detected by western blot after ALA and Tau exposure. D) Quantification of intensity of protein bands on western blot, normalized with the β-actin as a loading control.

### ALA enhances migration of microglia

Omega-3 fatty acid induces alternative anti-inflammatory phenotype of microglia; anti-inflammatory phenotype depicts increased migration of microglia. The alternative activation observed with IL-4 treatment to microglia induces excessive migration^35^. We hypothesize that exposure of ALA will modulate the cell membrane, and increased its fluidity, which showed the increased migration of N9 cells (Fig. 6a). In this study we checked for migration ability of microglia by wound scratch assay. Time-dependent migration of microglia was studied in presence of ALA for 0, 6, 12, and 24 hours (Fig. 6b). We quantified the number of cells into the wound after every time point by phase contrast microscopy images. ALA found to increase the migration of microglia as compared to cell control. At 24 hours’ time point monomer showed higher migration rate than aggregates, however with their respective exposure with ALA enhanced the migration to greater extent. The migration profile at 24 hours found to be highest in aggregates with ALA condition (Fig. 6c). These results suggest that ALA supports microglia to increase the migration, which is one of the key properties of anti-inflammatory phenotype of microglia.

**Figure 6.**
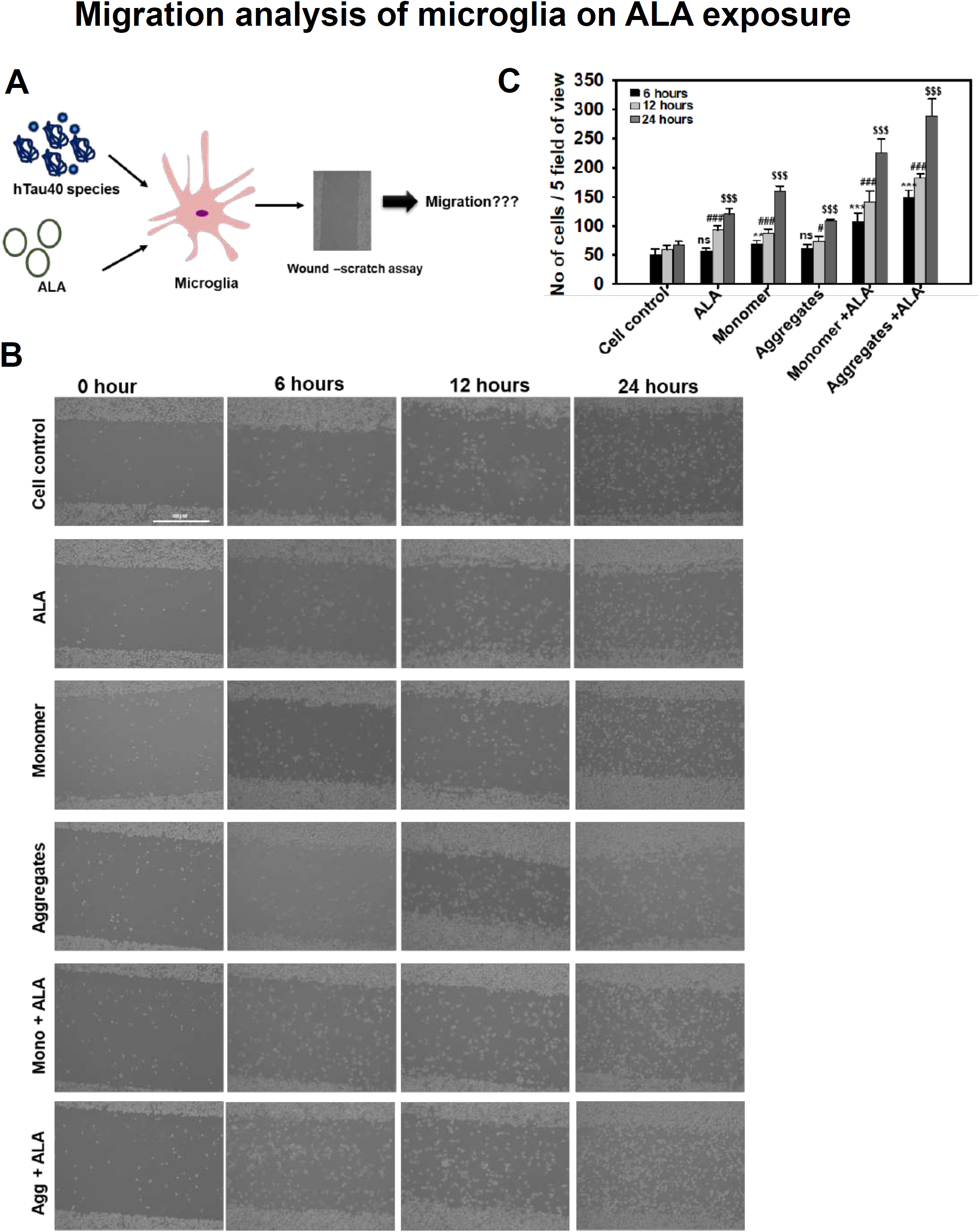
Migration analysis of microglia in presence of ALA. Increased migration microglia is a key property of anti-inflammatory phenotype. We desired to observe the effect of ALA on migration of microglia since omega-3 fatty acids enhance anti-inflammatory phenotype. A) ALA observed to enhance the phagocytosis of microglia that is also assisted by migration of microglia. The effect of ALA on migration under the influence of hTau40 monomer and aggregates has been studied. B) The migration of microglia was studied by wound scratch assay. Migration of cells into the scratch was studied with different time intervals 0, 6, 12, 24 hours after the scratch observed with phase contrast microscope. Scale bar is 100 µm. C) In each treatment groups random five fields were chosen and number of cells migrated into wound was counted. The comparison for each time point was carried out with its respective time point control (untreated) group; significance is P<0.001.

### ALA polarize nuclear-centrosomal axis in microglia

Increased migration is also associated with the repolarization of MTOC along the nucleus centrosome (NC) axis in the cell. In migratory cells the microtubule network reorient to migratory leading end for the forward motion and hence the MTOC positions are found predominately in the anterior region of nucleus. In case of highly migratory cells the repolarization of MTOC are observed to be present on different positions such as anterior, posterior and lateral positions to nucleus. The microtubule network found to be dense at the nucleus, spreaded towards lamellum and bundled down to uropod at rear end (Fig 7a). In case of ALA treated cells in both monomers and aggregates showed MTOC orientation in all the different positions, whereas in other treatment groups anterior position of MTOC prevails (Fig 7b). In case of aggregates with ALA treatment the percentage occurrence of different positions of MTOC around the nucleus is near to equal (Fig 7b). The picture representation suggests that ALA treatment to microglia reorient the MTOC to different positions anterior, posterior and lateral to nucleus unlike unipolar control cells (Fig 7c). All the panel of immunofluorescence images with DIC is shown indicating exact positions of MTOC (Fig. S3).

**Figure 7.**
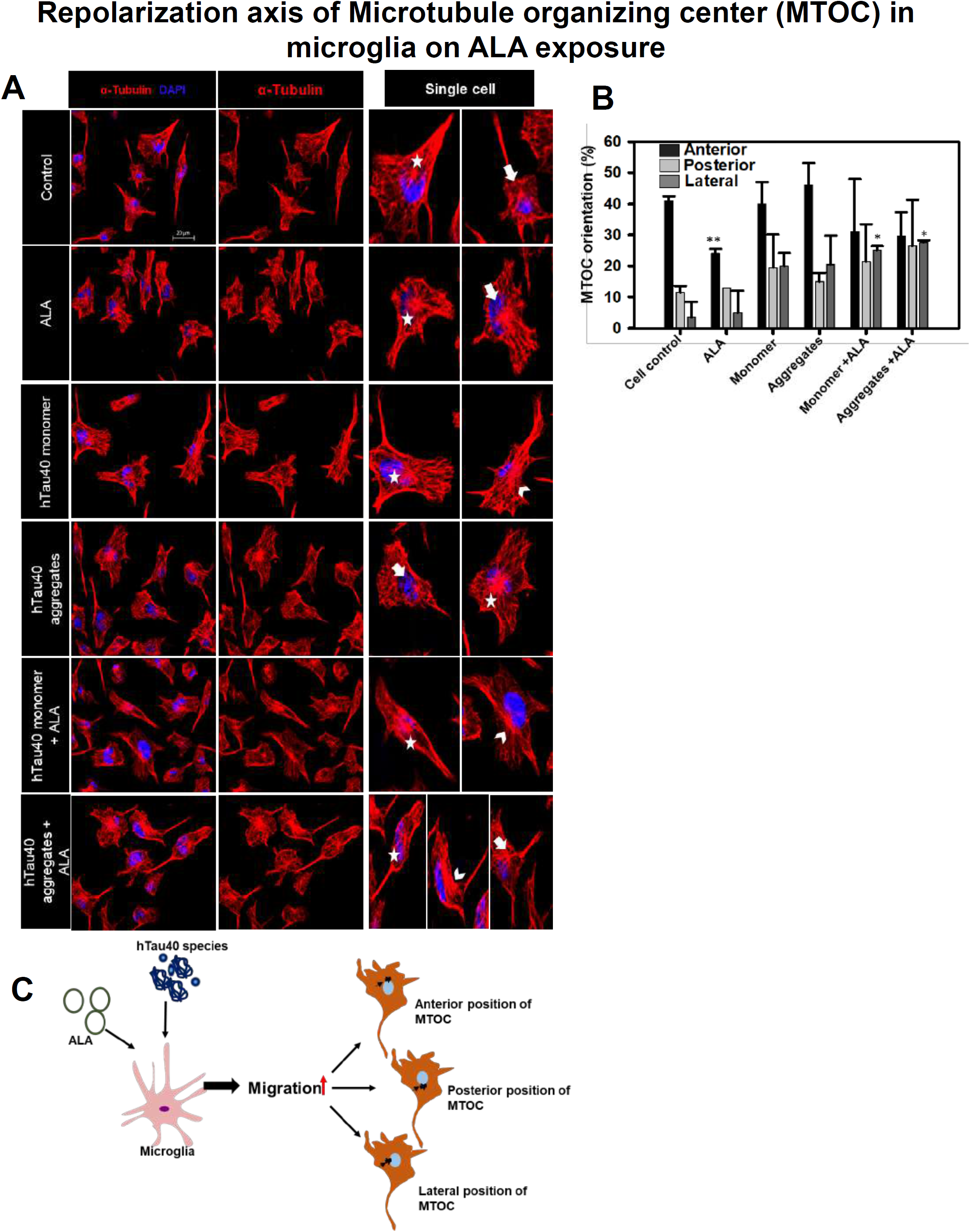
Repolarization of axis of MTOC on ALA exposure. A) Microglia were treated with hTau40 monomer, aggregates, ALA for 24 hours and observed for MTOC positioning with respect to nucleus in N9 cells. Fluorescence microscopy images were analyzed for MTOC positions stained with α-tubulin (red) and DAPI. The panels showing merge images and single filter images for tubulin are single cells with different MTOC position white star (anterior position), white open arrow head (lateral position), and white downward arrow (posterior position). Scale bar is 20 µm. B) Quantification of MTOC reorientation. The percentage number of cells for the different positions of MTOC was calculated in ten different fields in all the treated groups. C) Exposure of ALA with hTau40 monomer and aggregates found to increase the migration that modulates the orientation of MTOC positions in the cells. The expected different positions of MTOC are depicted according to pectoral representation.

**Figure 8.**
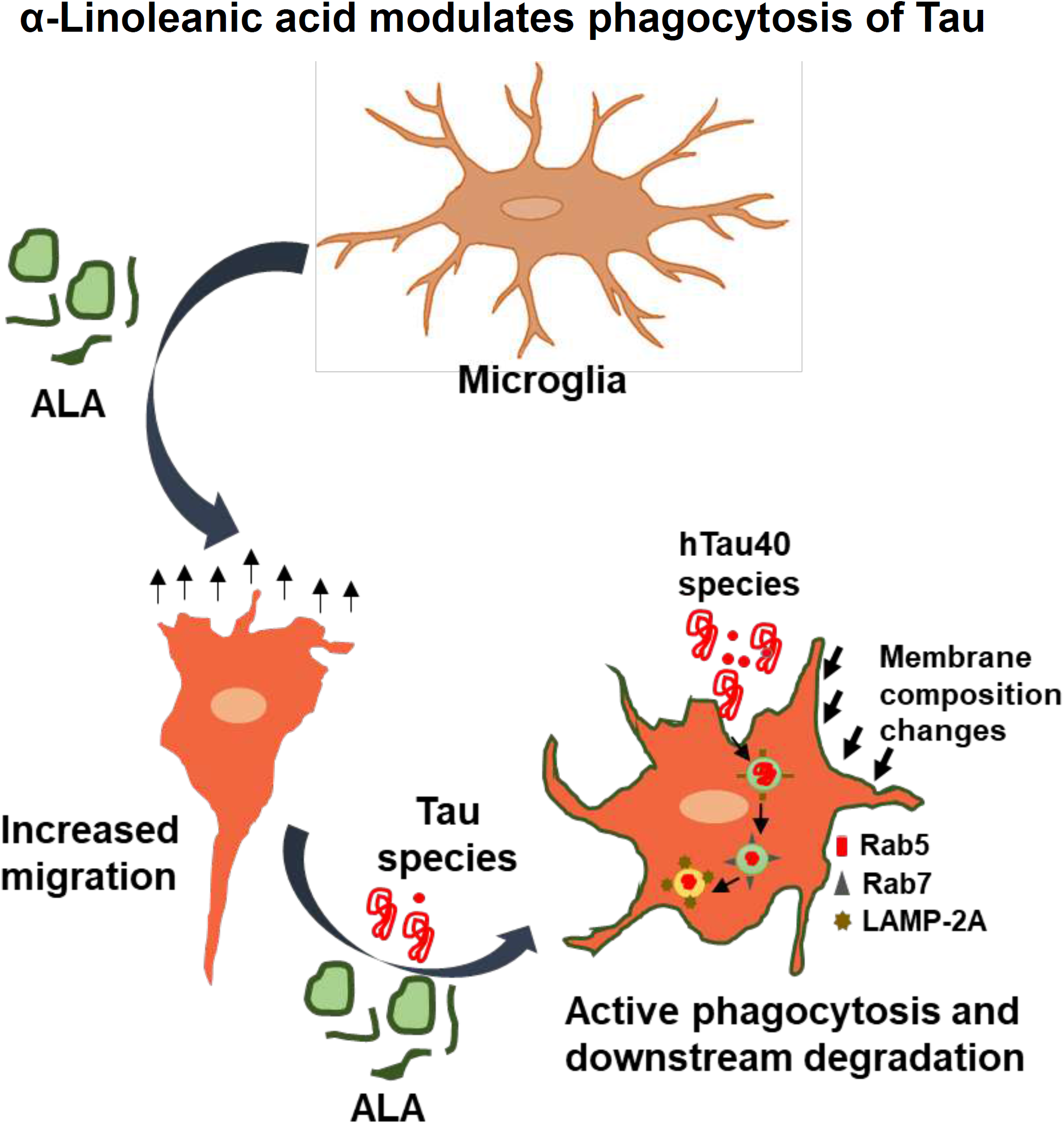
ALA modulates phagocytosis of Tau. ALA after exposure to N9 cells enhances extracellular Tau phagocytosis and its lysosome-mediated degradation. Thus reduces accumulation internalized Tau in microglia. ALA on the other hand enhances migration property of microglia to locate the target and undergo phagocytosis of extracellular Tau.

## Discussion

The extracellular Tau species after recognition by immune cells induces the immune response. Omega-3 fatty acidshas ability to effectively dampen the response given by immune cells. Dietary omega-3 fatty acids involves microglia into anti-inflammatory immune response, which would enhance clearance of extracellular pathological Tau species. The insoluble pathological aggregated form of Tau prepared *in vitro* was confirmed by SDS PAGE, TEM, ThS fluorescence and CD analysis. In present study, N9 microglia cells were exposed to ALA being an omega-3 fatty acid and observed for its beneficial effects. Exposure of ALA enhanced phagocytic ability of microglia by internalizing extracellular Tau species.The effect of ALA on migration has been studied on microglia cells as they assist the phagocytosis process. The enhanced phagocytosis in presence of ALA should also channel the internalized antigen towards lysosome-mediated degradation for desired clearance of extracellular antigens. The degradation of internalized Tau was denoted with the endosomal markers and their colocalization with internalized Tau. The reported results suggest the beneficial effects of ALA in brain.

In previous studies, it has been proven that the seeding nature of Tau as it causes template-dependent aggregation on uptake by healthy neurons ^11^. The aggregated extracellular Tau species secreted by various mechanisms have tendency to propagate the disease ^36-38^. The use of other omega-3 fatty acids DHA, EPA has been studied for the uptake of extracellular Aβ-plaques and their clearance ^14, 39^. To study the beneficial role of ALA for the uptake of extracellular Tau, we had incubated Tau and ALA with N9 microglia cells for 24 hours. The increased phagocytosis of extracellular Tau has been observed with ALA treatment conditions ^40^. The omega-3 fatty acids exerts ant-inflammatory properties to cells due to their ability to produce SPM (specialized pro-resolving molecules), which on attending certain concentration shows their effects ^41^. DHA and EPA are the main omega-3 fatty acids increase the microglial activation and act as a main precursor of SPM, they also found to activate PPAR-γ to mediate anti-inflammatory response ^13, 42^. The clearance of extracellular targeting species in case of AD could be objectified with dietary omega-3 fatty acids. Alzheimer’s disease is also being characterized by the endo-lysosomal abnormalities and accumulation of Rab5 positive enlarged endosomes followed by detectable Aβ-plaques ^16, 43^. The accumulation also impairs the fusion of autophagosomes with late-endosomes and lysosomal degradation. The transition of phagosomes from Rab5 to Rab7 is one of the important events, which specifies the degradation of internalized antigens ^44^. In our results the levels of Rab5 and Rab7 found to increase in case of ALA treatment and there is also significant colocalization of internalized Tau with Rab5 and 7 was observed indicating that internalized Tau is undergoing degradation pathway instead of accumulating inside the cell. The final step of phagosome maturation ends up with the fusion with lysosome. The formation of phagolysosome regulated by lysosomal-associated membrane proteins. Double knock out of LAMP-2A found to impairs the maturation of phagosome by halting the process prior to acquisition of Rab7 and affects lysosome density in cell ^34, 45^. LAMP-2A might be better target to study the lysosomal degradation of internalized Tau. We have observed from the results that ALA enhances LAMP-2A levels in cell and its colocalization with internalized Tau indicates the active phagocytosis.

Activation stage of microglia upon stimulation is observed with increased migration, phagocytosis, proliferation and cell shape changes which are assisted by actin cytoskeleton ^46^. The cell migration profile for N9 cells treated with different groups was studied by wound scratch assay. Excessive migration was seen with the ALA treatment as compared to control groups. The protrusive and contractile force needed for the migration is supported by actin rearrangements. The polarization of microglia is supported by both actin and microtubule cytoskeleton. In migratory polarized microglia well assisted reorientation of NC axis is observed, in many migratory cells anterior NC axis is observed where MTOC, endoplasmic reticulum and golgi apparatus are in front of nucleus which stabilized the front end. However in highly migratory ALA treated cells lacks the preference of NC axis and other positions such as posterior, lateral were observed ^35^. This is also observed in highly migratory immune cells such as neutrophils and T-lymphocytes. The increased migration supports enhanced phagocytosis in microglia.

## Conclusions

ALA enhanced phagocytosis of extracellular Tau as part of the anti-inflammatory property of microglia. The phagocytosis is coupled with the degradation of internalized Tau via lysosome-mediated degradation. This indicated beneficial role of dietary supplement of ALA over Tau seeding.

## Materials and methods

### Chemicals and Primary antibodies

Luria-Bertani broth (Himedia); Ampicillin, NaCl, Phenylmethylsulfonylfluoride (PMSF), MgCl2, APS, DMSO, Ethanol (Mol Bio grade), Isopropanol (Mol Bio grade) and methanol (Mol Bio grade) were purchased from MP biomedicals; IPTG and Dithiothreitol (DTT) from Calbiochem; MES, BES, SDS, α-Linolenic acid (ALA) (L2376) from Sigma; EGTA, Protease inhibitor cocktail, Tris base, 40% Acrylamide, TEMED from Invitrogen. For cell culture studies, the N9 microglial cell line no. is CVCL-0452, Roswell Park Memorial Institute (RPMI), Fetal Bovine Serum (FBS), Horse serum, Phosphate buffer saline (PBS, cell biology grade), Trypsin-EDTA, Penicillinstreptomycin, RIPA buffer were also purchased from Invitrogen. MTT reagent and TritonX-100, Trypan -Blue were purchased from Sigma. The coverslip of 12 mm was purchased from Bluestar for immunofluorescence and copper-coated carbon grids for TEM analysis were purchased from Ted Pella, Inc. In immunofluorescence and western blot study we used the following antibodies: Beta-actin (Thermofisher cat no. MA515739), Anti Alpha Tubulin Antibody Clone DM1A (Thermofisher cat no-62204), Tau Monoclonal antibody (T46) (Thermo cat no-136400), Anti-Iba-1 (Thermo cat no-PA527436), Rab5 (cell signaling, cat no 3547S), Rab7 (Santa Cruz Biotechnology, cat no Sc10767), LAMP-2A (Thermofisher cat no-51-2200), anti-mouse secondary antibody conjugated with Alexa Fluor-488 (Invitrogen, cat no A-11001), Goat anti-Rabbit IgG (H+L) Cross-Adsorbed Secondary Antibody with Alexa Fluor 555 (A-21428), GOXMS ALEXA FLOUR 488 goat anti rabbit (Thermofisher-cat no A28175) DAPI (Invitrogen), Goat Anti Mouse secondary antibody Peroxidase conjugated (Thermo fisher 32430), Prolong Diamond antifade (Thermofisher cat no-P36961).

### Protein expression and purification

Full-length wild type Tau protein (hTau40^wt^) was expressed in BL21* cells with 100 µg/ml of ampicillin antibiotic selection and purified with two-step chromatography methods, cation-exchange chromatography and size-exclusion chromatography (Gorantla, MiMB, 2018). Cells were grown at 37°C, scaled up and harvested after induction with 0.5 mM IPTG for 4 hours. Cells were subjected to homogenization to produce cell lysate at 15000-psi pressure. The cell lysate was subjected to 90°C heating in presence of 0.5 M NaCl and 5mM DTT for 20 min to denature structured proteins. The supernatant was collected after centrifugation at 40000 rpm for 45 minutes then put through dialysis overnight in 20 mM MES buffer supplemented with 50 mM NaCl. The supernatant was obtained again after centrifugation at 40000 rpm for 45 min passed through cation-exchange chromatography. Sepharose fast-flow column was used for chromatography, using 20 mM MES buffer and 50 mM NaCl (Buffer A). Elution was done with 20 mM MES buffer and 1 M NaCl (Buffer B). A fractions containing Tau proteins were collected after cation exchange chromatography, it was then concentrated and subjected to size-exclusion chromatography. Size-exclusion chromatography was carried out in the Superdex 75 Hi-load 16/600 column in 1X PBS supplemented with 2 mM DTT. A fractions containing Tau were collected, pooled, concentrated and the concentration of protein was determined with BCA (Bicinchoninic acid assay) assay.

### Aggregation assay

Tau protein undergoes aggregation in presence of poly-anionic reagent such as heparin and arachidonic acid, etc., it is observed by the transition of random coiled structure to the β-sheet formation in protein ^25^. In this study Tau aggregation was induced by heparin (MW-17500 Da) in the ratio of 1:4 heparin to Tau along with other additives 20 mM BES buffer, 25 mM NaCl, 1 mM DTT, 0.01% NaN_3,_ PIC. The effect of ALA on Tau aggregation was measured by Thioflavin S (ThS) fluorescence assay. ThS is a homogeneous mixture of methylation product of dehydrothiotoluidine in sulfonic acid, which can bind to β-sheet structure. Aggregation kinetics of Tau was studied with 2 µM of Tau and ThS in 1:4 ratios. The excitation wavelength for ThS is 440 nm and the emission wavelength is 521 nm, further analysis of data was done using Sigmaplot 10.0.

### Transmission electron microscopy

Morphological analysis of Tau fibrils and ALA vesicles were studied by transmission electron microscopy (TEM). 2 µM Tau sample was incubated on 400 mesh, carbon-coated copper grid and stained with 2% uranyl acetate. For ALA vesicles working concentration of 40µM was taken for grid preparation. The images were taken with TECNAI T20 120 KV.

### CD spectroscopy

Conformational changes in Tau from random coiled structure to β-sheet conformation on aggregation of protein was studied using CD spectroscopy, the spectra was collected as previously mentioned in UV region ^26^. The measurement was done in Jasco J-815 spectrometer, cuvette path length was 1 mm, measurement was done in range of 250 to 190 nm, and with a data pitch of 1.0 nm, and scanning speed was kept 100 nm/min. For measurement 3 µM sample concentration was taken in phosphate buffer pH 6.8 all the spectra were taken at 25°C.

### Cell culture and preparation of ALA

N9 (microglia) cells were grown in RPMI media in T25 flask or 60mm dish supplemented with 10% heat-inactivated serum, 1X penicillin-streptomycin antibiotic solution and glutamine for maintain the culture. Cells were passaged on 90% confluence using 0.25% trypsin-EDTA solution after washing with PBS. For western blotting experiment cells were seeded in 6 well plate. For α-Linolenic acid preparation, previously published protocol was followed ^14^. Briefly, ALA was dissolved in 100% molecular biology grade ethanol and solubilized at 50°C in the stock concentration of 20 mM The fatty acid solution was prepared freshly before every experiment. According to the previous studies, 40 µM was the working concentration of ALA for carrying further experiments. The final concentration of ethanol in cell culture media was maintained below 0.5%.

### Tau internalization

To study the effect of ALA on microglial phagocytosis, N9 cells were treated with extracellular 1 µM monomer and aggregates along with 40 µM ALA. For the immunofluorescence experiment (25,000 cells/well), N9 cells were seeded on 12 mm glass coverslip in 24 well-plate. The cells were then incubated with 1 µM Tau monomer and aggregates along with 40 µM ALA for 24 hours. To compare the internalization ability the controls of 1 µM Tau monomer and aggregates alone for the comparative studies with ALA treatment, 40 µM ALA alone to check morphological changes in N9 cells and cell control (without treatment) were kept. The coverslips were then fixed and stained for immunofluorescence analysis with antibodies Tau T-46 (1:400 dilution) and Iba-1 (1:500 dilution). The mounting of coverslips were done with mounting media (80% glycerol in 1X PBS). The intracellular intensity of microglia were calculated from fluorescence images to quantify the internalization. The representation of intracellular intensity was done as an intensity/µm area.

To study the degradation of intracellular Tau after phagocytosis we have targeted early and late endosomal markers and lysosome marker for final degradation process. The treatment was done as previously mentioned, after 24 hours of exposure cells were fixed and stained for immunofluorescence analysis. The analysis of the process of degradation was done co-localization of internalized Tau with Rab 5 (1:200 dilution), Rab7 (1:200 dilution) and LAMP-2A (1:500 dilution). The intracellular intensity of Rab 5, 7 and LAMP-2A were studied as intensity/ µm area to understand the expression of proteins on ALA exposure. The colocalization of internalized Tau was studied with 3-D and orthogonal analysis of immunofluorescence images.

### Wound scratch assay

To study the migration of microglia wound-scratch assay was performed. For the assay, (5,00,000 cells/well) N9 cells were seeded in a 6-well plate and maintained in RPMI media for 24 hours till the confluency reached to 80%. Scratch was created with sterile 200 µl pipette tip, followed by treatment with groups as mentioned previously. Cells were incubated further for 24 hours to study the migration of N9 cells into the wound. A number of cells migrated into the wound were calculated for 5 different areas of culture and the average was calculated to quantify the migration.

### MTOC reorientation analysis

To study immunofluorescence experiment (25,000 cells/well) N9 cells were seeded on 12 mm coverslips in 24-well plate. The desired treatment of Tau monomer, aggregates and ALA was given to cells for 24 hours and fixed for immunofluorescence staining. The MTOC positions were analyzed by α-tubulin (1:200) staining. The anterior, posterior and lateral positions of MTOC were counted with respect to the nucleus denoted by DAPI stain. The percentage of MTOC positions were calculated in 10 different fields.

### Immunofluorescence analysis

N9 cells were passaged in RPMI media supplemented with 10% FBS and 1X penicillin-streptomycin. For immunofluorescence studies, 25,000 cells were seeded on 12 mm coverslip (Bluestar) in 24 well plate. Supplemented with 0.5% serum-deprived RPMI media for the desired treatment. The treatment was given for 24 hours. Cells were then fixed with chilled absolute distilled methanol for 20 minutes at -20°C then washed with 1X PBS thrice. Permeabilisation was carried out using 0.2% Triton X-100 for 15 Minutes, washed three times with 1X PBS followed by blocking with 2% serum in 1X PBS for 1 hour at room temperature. Primary antibody treatment was given to cells overnight at 4°C in 2% serum in 1X PBS in a moist chamber. The next day, cells were washed with PBS thrice. Then incubated in the desired secondary antibody in 2% serum at 37°C for 1 hour. Further cells were washed with 1X PBS 3 times and counterstained with DAPI (300 nM). Mounting of coverslip was done in mounting media (80% glycerol). Images were observed under a 63x oil immersion lens in Axio observer 7.0 Apotome 2.0 Zeiss microscope.

### Western blot

For detection of protein levels in cells (3,00,000 cells/well) N9 Cells were seeded in 6 well plate and after the desired treatment for 24 hours. Treatment exposure followed by washing with 1X PBS. Cell lysis was carried out using radioimmunoprecipitation (RIPA) assay buffer containing protease inhibitors for 20 min at 4°C. The cell lysate was centrifuged at 12000 rpm for 20 minutes. Protein concentration was checked by using Bradford’s assay and equal amount of 75 µg total proteins for all the treatment groups were loaded on polyacrylamide gel electrophoresis of range 4-20% and the gel is electrophoretically transferred to polyvinylidene difluoride membrane and kept for primary antibody Rab5, Rab7, LAMP-2A, Iba-1 (1:1000 dilution) binding for overnight at 4°C. After the incubation washing of blot was carried out three times with 1X PBST (0.1% Tween-20). The secondary antibody were incubated for 1 hour at RT. Then the membrane was developed using chemiluminiscence detection system. The relative quantification of protein was carried out with loading control β-Actin (1:5000) in each treatment group.

### Statistical analysis

All the experiments have performed 3 times. The data is analyzed using SigmaPlot 10.0 and the statistical significance was calculated by student’s *t*-test (ns-non-significant, * indicates P≤0.05, ** indicates P≤ 0.01, *** indicates P≤0.001). The quantification of levels of intracellular proteins in immunofluorescence experiments was carried out by measuring the absolute intensity of protein and the corresponding area of microglia with Zeiss ZEN 2.3 software for image processing.

## Abbreviations

ALA: α- Linolenic acid
CNS: central nervous system AD- Alzheimer’s disease
PUFAs: polyunsaturated fatty acids
DHA: Docosahexaenoic acid
EPA: Eicosapentaenoic acid
NFTs: Neurofibrillary tangles
MTOC: microtubule organizing center
NC axis: nuclear-centrosomal axis.
LAMP-2A: lysosome associated membrane protein

## Acknowledgements

We are grateful to Chinnathambi’s lab members for their scientific discussions, helpful suggestions and we highly appreciate critical reading of the manuscript. We are thankful to Dr. Deepa Subramaniyam NCCS, Pune to provide antibodies for endosomal-pathway analysis. Authors

## Authors’ contributions

SD and SC performed the experiments and prepared the initial draft. SC conceived, designed, supervised, initial draft, review editing and wrote the paper. All authors read and approved the final paper.

## Author’s information

**Smita Eknath Desale**

*Neurobiology Group, Division of Biochemical Sciences, CSIR-National Chemical Laboratory, Dr. Homi Bhabha Road, 411008 Pune, India*

**Smita Eknath Desale**

*Academy of Scientific and Innovative Research (AcSIR), 411008 Pune, India*

**Subashchandrabose Chinnathambi**

*Neurobiology Group, Division of Biochemical Sciences, CSIR-National Chemical Laboratory, Dr. Homi Bhabha Road, 411008 Pune, India*

**Subashchandrabose Chinnathambi**

*Academy of Scientific and Innovative Research (AcSIR), 411008 Pune, India*

## Corresponding author

Correspondence to Subashchandrabose Chinnathambi

## Supplementary information

**Figure S1.**
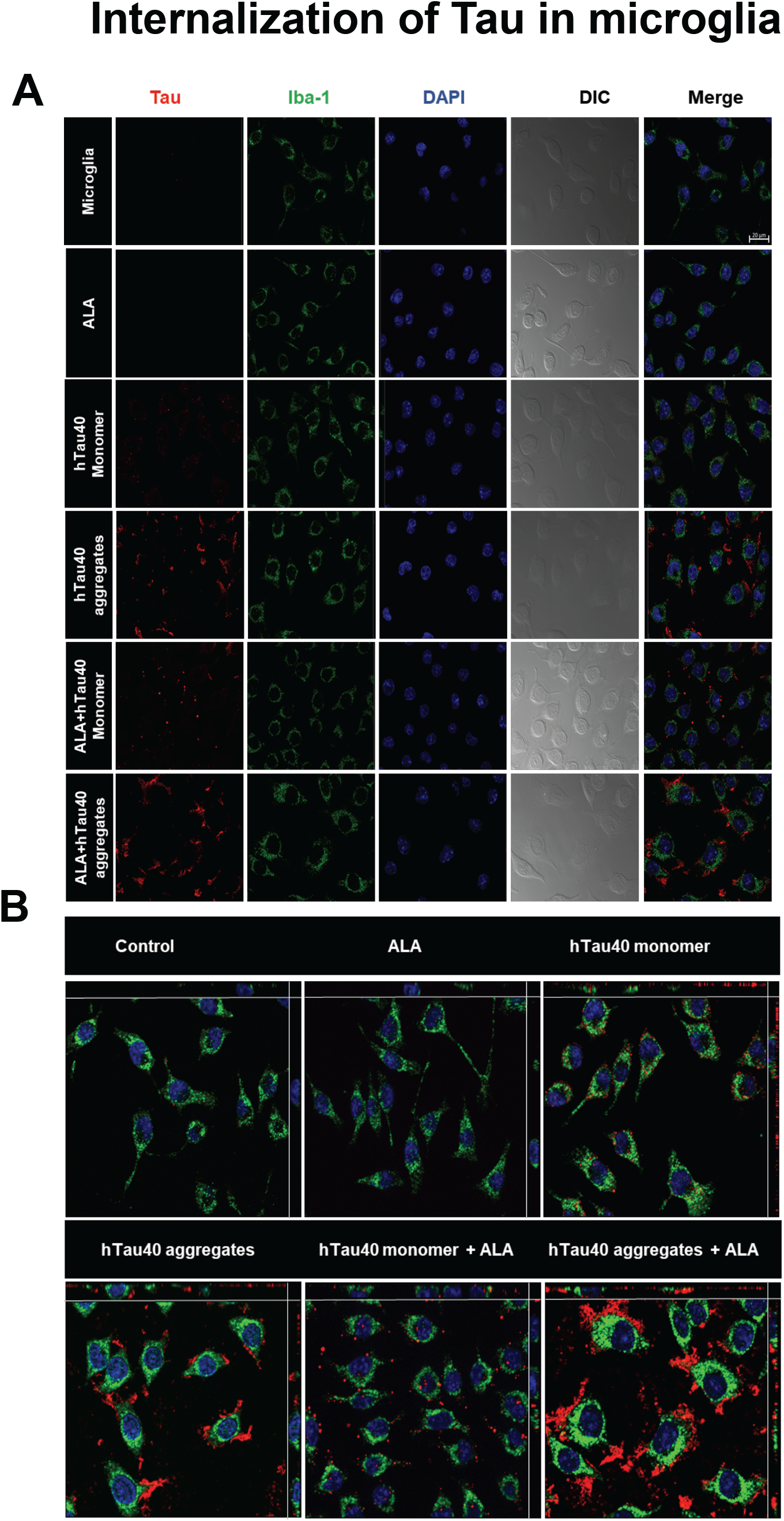
Internalization of extracellular Tau in microglia. Internalization of extracellular Tau in microglia induced by ALA. N9 cells were treated with extracellular Tau monomer and aggregates along with the ALA. After 24 hours of treatment cells were stained for Tau (red), Iba-1 (green), DAPI. A) Individual panel of 2D immunofluorescence images of Tau internalization. Panel contains Tau, Iba-1 DAPI, DIC and merge images. B) Orthogonal view of immunofluorescence images of Tau internalization. The images are represented with x and y-axis indicating the position of Tau in microglia. Scale bar is 20 µm.

**Figure S2.**
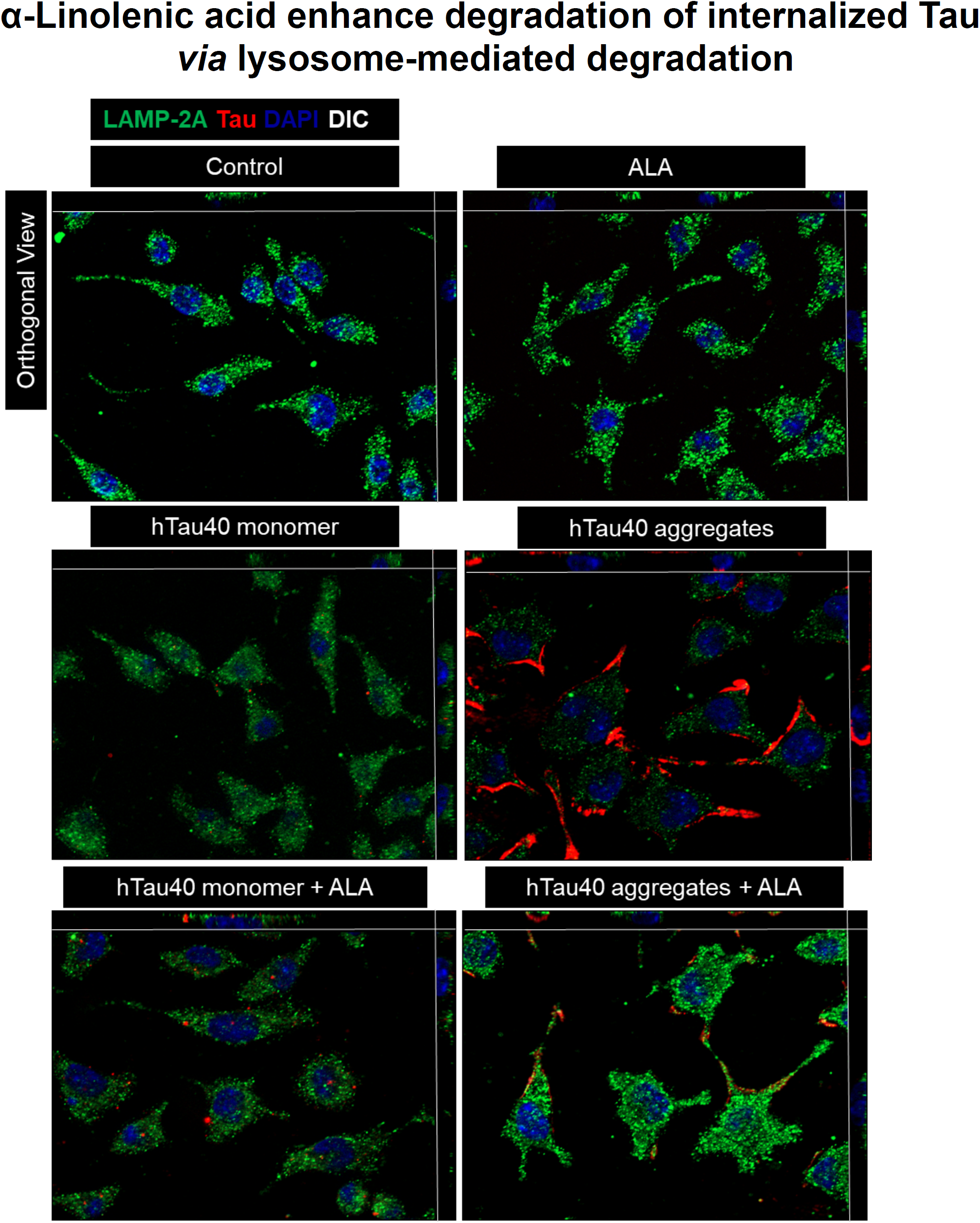
α-Linolenic acids enhance degradation of internalized Tau via lysosome-mediated degradation. Internalization of extracellular Tau in microglia induced by ALA was studied for the lysosome mediated degradation via LAMP-2A lysosomal marker. N9 cells were treated with Tau monomer aggregates and ALA and stained for LAMP-2A (green), Tau (red) and DAPI. A) Immunofluorescence images showing orthogonal view of the merge image of Tau and LAMP-2A staining. The x and y axis of the orthogonal view indicates position of Tau and its colocalization with LAMP-2A. Scale bar is 20 µm.

**Figure S3.**
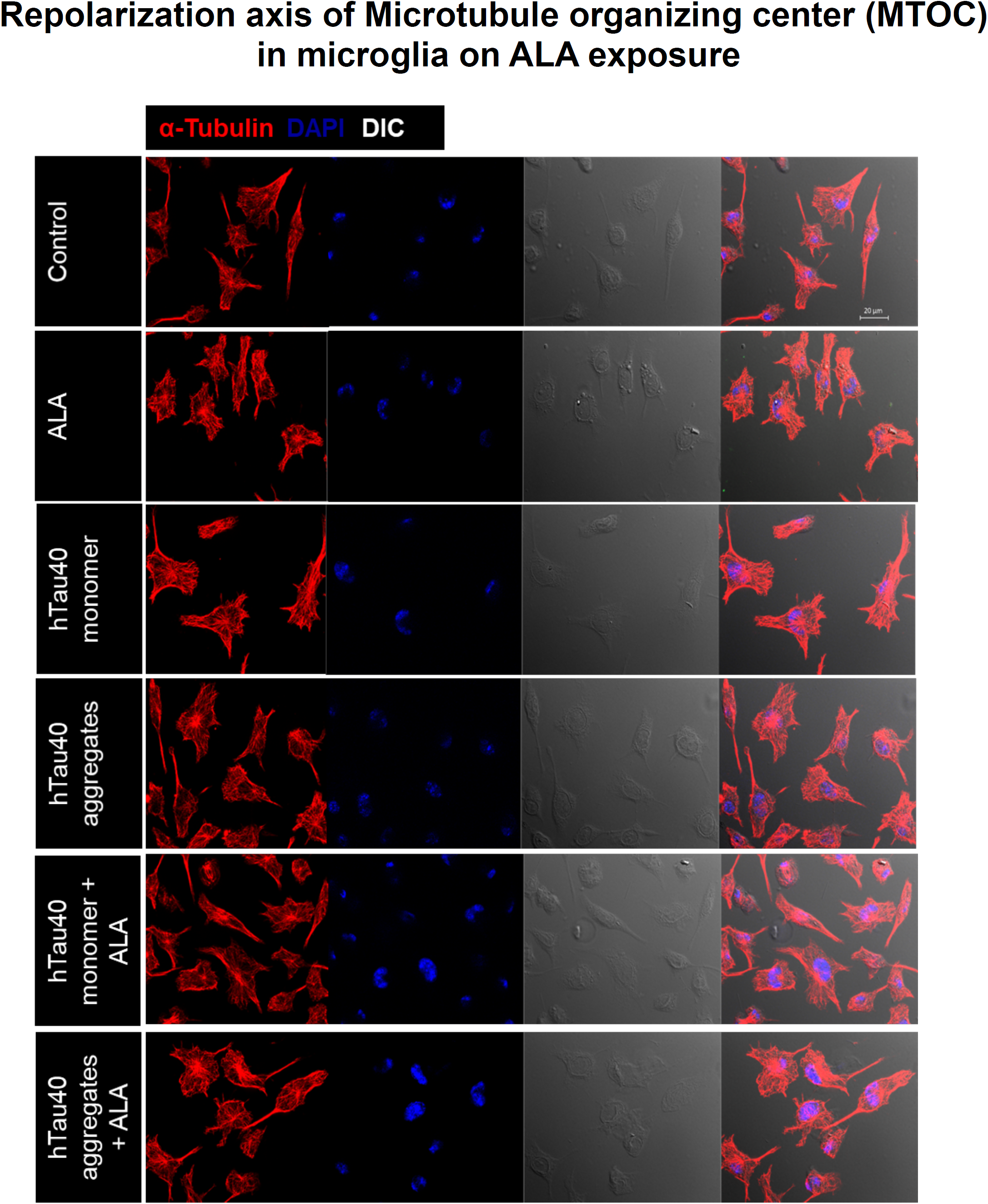
Repolarization axis of microtubule organizing center (MTOC) in microglia on ALA exposure. Polarization of axis of MTOC in microglia was studied as one of the property of migratory cells. Repolarizations of MTOC positions were studied upon ALA exposure. A) Immunofluorescence images showing separate panel for microtubule (red), DAPI and DIC. Merge image with DIC indicates positions of MTOC in cell. Scale bar is 20 µm.

